# TReNCo: Topologically associating domain (TAD) aware regulatory network construction

**DOI:** 10.1101/2021.04.27.441672

**Authors:** Christopher Bennett, Viren Amin, Daehwan Kim, Murat Cobanoglu, Venkat Malladi

## Abstract

There has long been a desire to understand, describe, and model gene regulatory networks controlling numerous biologically meaningful processes like differentiation. Despite many notable improvements to models over the years, many models do not accurately capture subtle biological and chemical characteristics of the cell such as high-order chromatin domains of the chromosomes. Topologically Associated Domains (TAD) are one of these genomic regions that are enriched for contacts within themselves. Here we present TAD-aware Regulatory Network Construction or TReNCo, a memory-lean method utilizing epigenetic marks of enhancer and promoter activity, and gene expression to create context-specific transcription factor-gene regulatory networks. TReNCo utilizes common assay’s, ChIP-seq, RNA-seq, and TAD boundaries as a hard cutoff, instead of distance based, to efficiently create context-specific TF-gene regulatory networks. We used TReNCo to define the enhancer landscape and identify transcription factors (TFs) that drive the cardiac development of the mouse. Our results show that we are able to build specialized adjacency regulatory network graphs containing biologically relevant connections and time dependent dynamics.

## Introduction

It is of critical importance to understand, model, and describe gene regulatory networks (GRN) that control diverse cellular functions of interest like those that drive differentiation or transitions from one development stage to another (Lee et al. 2002; DeRisi et al. 1997; Goode et al. 2016). With the advent of next generation sequencing technologies, it is now commonplace to reconstruct these networks to connect transcription factors (TFs) to the genes they regulate (Karlebach and Shamir 2008). One classic method is integration of cis-regulatory elements, like enhancers, and gene expression via matrix factorization to form network graphs between genes and TFs (Marbach et al. 2016). Generally, this is done using Chromatin Immunoprecipitation (ChIP) for H3K27ac to identify enhancers and RNA-seq to identify controlled genes. In many cases connections are determined through perturbations in upstream components like TFs and observing resultant changes in downstream expression levels (Gasperini et al. 2019). This method works exceptionally well for certain classes of TF and for closely linked enhancer-gene interactions. However, it commonly uses arbitrary length cut offs to prevent enhancers from erroneously influencing genes in distant parts of the genome. This can lead to enhancers having shorter or broader ranges of influence than what occurs biologically. As many recent chromosome-confirmation-capture (e.g. 5C, Hi-C and ChIA-PET) experiments have shown, there can be very broad and dynamic interactions made between different parts of a chromosome (Branco and Pombo 2007; McCord et al. 2020). Thus, it is more relevant to dynamically limit enhancers range of influence to only the topologically linked portions of the genome an enhancer is confined to, also known as Topologically Associated Domains (TADs). These regions are highly conserved across cell types and are known to limit the influence of cis-regulatory elements by physically separating them (Beagan and Phillips-Cremins 2020). Thus, it is critical that these cutoffs are included in the model to fully represent and capture the true biological processes occurring.

Here we present TAD-aware Regulatory Network Construction or TReNCo, a powerful, memory efficient tool for constructing regulatory networks from enhancer, promoter, and gene expression data without the need for perturbations. We designed TReNCo to construct a graph of interaction weights between TFs and the genes that they control using TAD boundaries to dynamically limit the range of enhancer influence. We utilize dynamic programming to factor matrices within TADs and combine network into a full adjacency matrix for a regulatory graph. With this method, we are able to capture biologically relevant interactions between known TFs and their gene targets. We show that this network contains many subtle interactions that could be a treasure trove of novel or uncharacterized interactions. We believe this method opens the possibility for understanding deeper mechanistic connections and new possibilities for identifying biological targets for drug discovery.

## Results and Discussion

### Model Design and Function

We designed TReNCo utilizing a previously reported core matrix factorization method with a distance-based scoring system broken down into subunits based on TAD boundaries (Marbach et al. 2016; Cuellar-Partida et al. 2012). In brief, our algorithm uses normalized gene expression count tables from RNA-seq (tsv files) and H3K27ac ChIP-seq read alignments (bam files), peaks (bed files), and TAD boundaries (a bed file) (Figure 1). Though the source of these data can vary, we designed TReNCo with ENCODE uniform processing pipelines in mind. We first generated initial expression matrices for gene expression (G) and enhancer expression (E) by sample. This was accomplished using the count tables from RNA-seq and building count tables for all enhancers merged into non-overlapping segments from the ChIP-seq data. These counts were used to calculate the Transcripts/Fragments per Kilobase Million (TPM) which are then log-scaled. A key file, provided by the user, is used to link related files to build a full expression matrix and, secondarily, serves to reduce memory usage by allowing batch processing of data.

**Figure 1:**
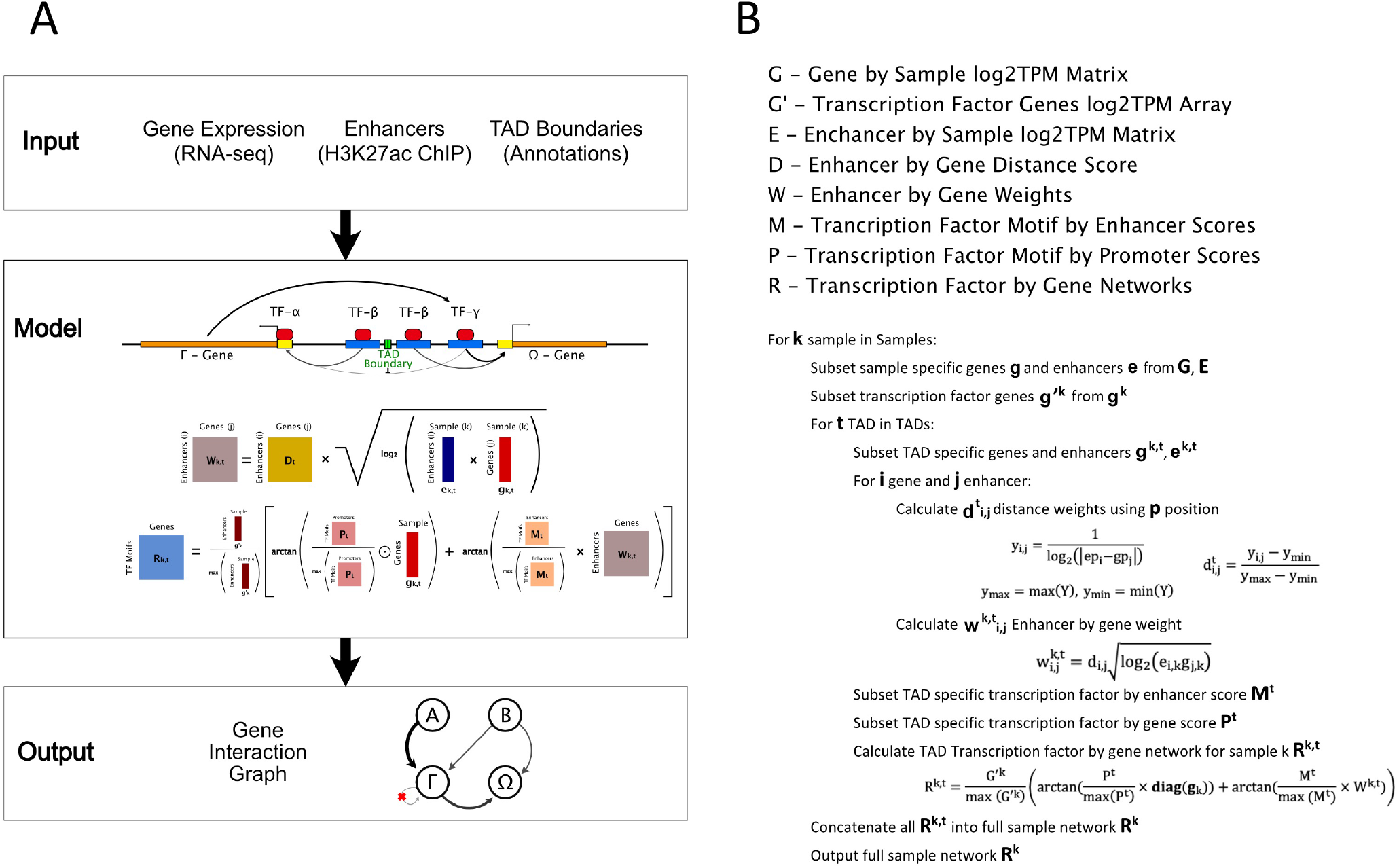
TReNCo model. A) Diagram with required inputs, gene expression, enhancers, and TAD boundaries input into a model using the basic equations shown leading to a gene interaction graph result. ‘+’ and ‘x’ indicate standard matrix additions and multiplications, respectively. The other operations such as ⊙, log, square root, and arctan are all element-wise. B) Pseudo-code for constructing a Gene Interaction Graph.

We next worked to establish TF-gene linkages by identifying TF binding sites in promoters and enhancers using a well-known program, FIMO (a part of the MEME software suite and report the log-odds score of TF binding) a major weight needed for establishing interaction (Grant et al. 2011). We designed a simple pipeline to generate promoter and enhancer master bed files and remove any potential overlaps between promoters and enhancers to ensure that TFs are not double counted to a gene. Furthermore, these files contained the union of all promoters and enhancers between the samples in order to streamline the identification of TFs. This was a critical time saving step as FIMO cannot be multithreaded and can take upward of 24 hours to run. By using master files, we could run FIMO only once per process leading to a huge performance boost.

With these core datasets, for each sample we were able to select sample-specific genes, enhancers, and TF interactions. To ensure proper TAD boundaries were followed and to improve speeds through multithreading, we designed a dynamic programming algorithm to process these datasets by TADs and generate TAD-specific distance weight matrices (D^t^) for each set. These matrix subsets were factored with the square-root of a TAD-specific interaction matrix produced via vector multiplication between the gene (g^k,t^) and enhancer (e^k,t^) expression profiles resulting in a TAD-weight matrix (W^k,t^). To generate an enhancer-specific graph edges, the weight matrix was factored with the TAD-specific enhancer-TF by gene matrix (M^t^) normalized to the maximum value of the matrix. This was done to set a standard scale of log-odds that was comparable between enhancers and promoters. We designed this component with the assumption that enhancer-TF binding should be similar in promoters and should be weighted the as a log-odds scale in the network. A promoter-TF by gene specific subnetwork (P_t_) was produced in a similar manner as the enhancer-specific network with weighing done using a TAD-specific gene expression vector since all distances between promoters and genes are 1. Arctangents were applied to both matrices due to the properties of the transformation where larger values approach an asymptote of π/2 while smaller values are approximately scaled linearly. This scaling draws larger value outliers into a tighter range without heavily influencing lower values and assumes a maximum impact a TF can have on a gene. The resulting matrices were added together and further weighted by normalized TF gene expression to lower the influence of lowly expressed TFs while minimally changing the effects from highly expressed TFs. The resulting TAD subgraphs were concatenated together into a full network adjacency graph matrix. Since this was a sparse matrix, TReNCo represents it as an adjacency array allowing us to store the information in much less space than is needed for a matrix.

### Model Validation

To validate the model, we used the extensive cohort of matched gene expression and H3K27ac ChIP-seq analyses in ENCODE (Davis et al. 2018) (Table S1). We decided to use mouse heart data due to the abundance of well correlated time point data spanning embryonic day 10.5 to 8 weeks after birth, highly characterized heart developmental processes, and the availability of previously documented TF-gene networks (Akerberg et al. 2019; Schlesinger et al. 2011) (Figure 2). While the ChIP-seq data is not highly correlated across all the sample types, the gene expression data has an R-squared of at least 0.7 between different biological samples. One e14.5 experiment set had an average R-squared of approximately 0.6 with all other biological samples. To remove this potentially problematic dataset in this analysis before the larger more computationally expensive processes occur, we added an optional soft filter in TReNCo to automatically remove any samples with an average R-squared less than 0.7 across all samples, for gene expression data. We were left with a set of highly correlated data that led us to conclude that this dataset was sufficient to use to TReNCo.

**Figure 2:**
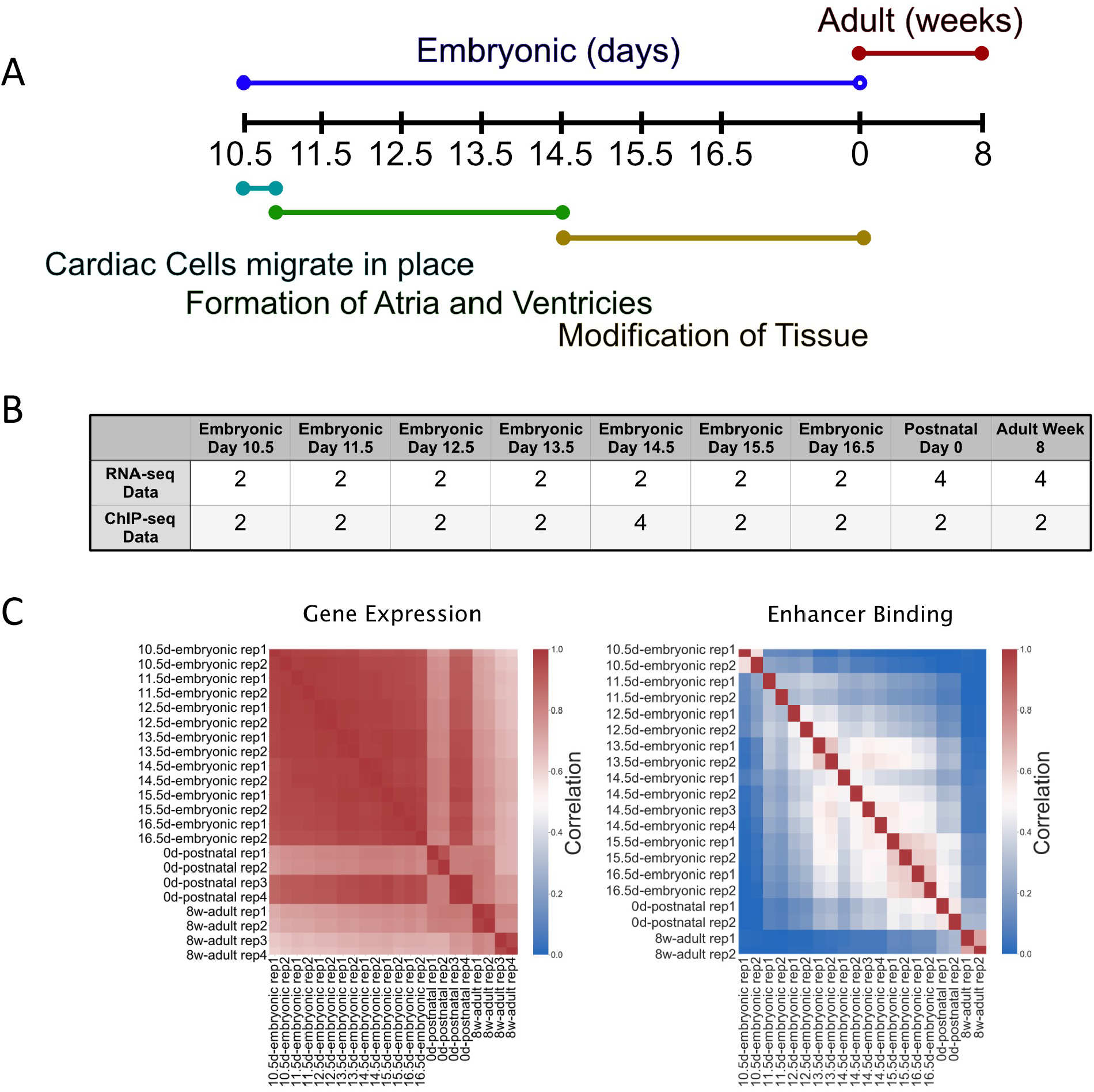
ENCODE Datasets. A) Timeline of basic mouse cardiovascular development with life stage on top and developmental stages on the bottom B) Number of samples for each timepoint and data type C) Correlation heatmap between samples and replicates at each time point

Previous studies of the mouse heart have identified Gata4, Mef2a, Nkx2-5, Tbx5, and Srf as important embryonic lethal TFs critical for development (Gittenberger-De Groot et al. 2005). When looking at the distribution of these TFs over time, we observed that there are many subtle dynamics in how the TFs’ weights shift. Gata4, Mef2a, Tbx5, and Nkx2-5 show a multimodal distribution with three major peaks and varying differences between time points though mostly the distributions overlapped (Figure 3A, Figures S2, S3, S4). We found that the weight distribution followed a similar trend; the dominant population of edge weights appears less than 0.1, a second mid population is between 0.1 and 0.3, and a final population above 0.3 that stretches up to 1. Another TF, Foxs1, demonstrated a more pronounced time point dependent change in addition to a tri-modal edge score (Figure 3B). Interestingly, Srf did not show this trend and tended to have lower weight edges throughout the distributions. To visualize the timepoint dynamics more clearly, we generated a heatmap of the distributions with inflection points added to determine changes in gene weights that may occur (Figure 3C and D). Inflection points, in this case, are simple differences in weights between each time point and the previous time point. These data are ideal for highlighting changes between each time point. An additional differential heatmap of all weight differences with respect to the embryonic day 10.5 point was generated to visualize change from a central time (Figure S1). It was clear that these TFs have time dependent dynamics in our model. Gata4, Nkx2-5, and Tbx5 appear to interact with most of their targets constantly throughout early development as indicated by a mostly yellow (no change) inflection point heat map until adult heart. These TF’s have been shown to be important in normal cardiac development (Misra et al. 2014) and act as potential as cardiac reprogramming factors from embryonic fibroblast (Hashimoto et al. 2019). At this time, we observed a net decrease in the Gata4 network weights as observed by an increase in negative inflection points and a decrease in positive values. Mef2a showed a similar trend as Gata4 with a minor increase in network weights leading up to birth, which has been previously shown to be important in postnatal heart development and regulation (Desjardins and Naya 2016). Srf shows a different trend with most of the weights being relatively low until P0 where there is a minor but noticeable uptick in the network weights. This observation matches the biological importance of Srf in early cardiac development and its critical role in maintaining adult heart function (Mokalled et al. 2015). Foxs1 demonstrates the most profound change over time with the initial weights being very low and increasing over time until embryonic day 16.5. After this time the weights begin to decrease into adulthood but never go away completely. This may be due to the role of Foxs1 as a key factor in vascular development (De Val 2011) which in important in earlier development.

**Figure 3:**
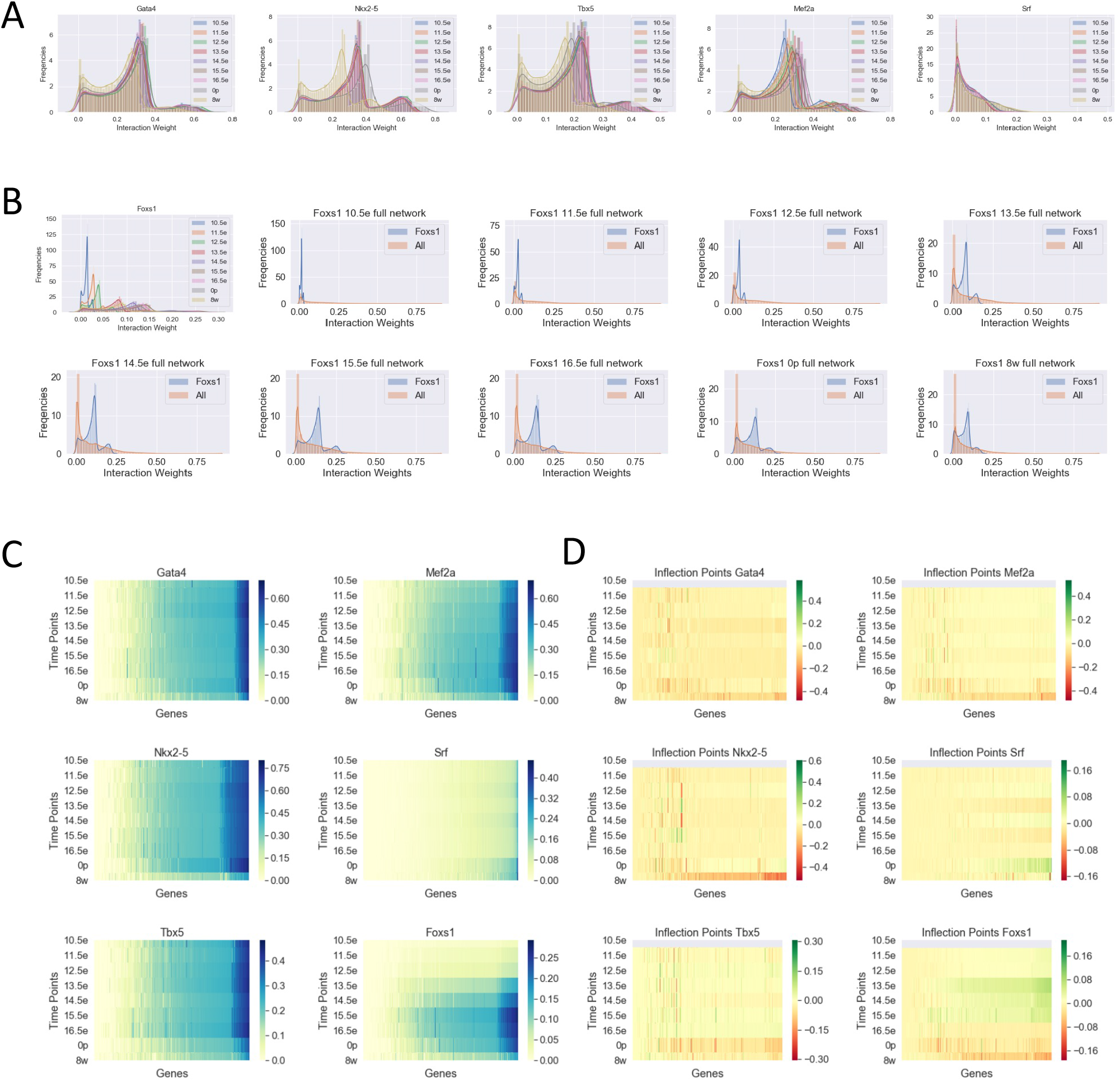
TF-Gene Edge Weights. A) Histograms of TF-gene interaction weights for 5 different genes, separated by developmental time points B) Histogram as in A, for Foxs1 TF separated into individual developmental time point plots C) Heatmap of TF-gene interaction weights sorted by time points D) Heatmap as in C, showing gene inflection points calculated by log2 the ratio of gene weights. Green indicates increase gene weight from the last time point while Red indicates a decrease.

### Model Comparison

There have been a number of studies on mouse cardiac TF regulatory networks with one study looking at the regulatory networks of Gata4, Mef2a, Nkx2-5, and Srf and providing the interactions they identified (Schlesinger et al. 2011). We extracted the interactions of the aforementioned TFs from our network and compared it with the previously identified interactors (Figure 4A and S8). We found that our networks contain over 10,000 putative novel interactions (weight edge weight greater than 0) that were not reported previously. Interestingly, regardless of the timepoint, our networks captured only about 63% of Gata4 targets, 61.5% of Mef2a targets, and 57% Nkx2-5 and Srf targets of the previous network’s interactions leaving a large portion of their networks unique to their analysis (Figure 4B). We speculate there are two likely explanations for the absence of a 100% overlap: 1) the previous network established interactions using the canonical distance-based cutoff leading to some genes being added or removed erroneously if cutoffs differed from our TAD boundaries or 2) while our TAD boundaries are more accurate than distance-based cutoffs, the TADs we use are not fully representative of cardiac specific TADs leading to loss of some connections in our network. Regardless of the reason, we wanted to understand if the main overlap between our networks was due to the previous study finding the strongest interactors of the TFs. To test this, we performed the Kolmogorov–Smirnov test (KS-test) on the cumulative distribution between the overlap edge weights and the full edge weights (Figure 4C and Table S2). We found that in all time points the overlapping genes identified have higher mean edge weights than the total (Figure 4D). This implies that we are identifying true strongly interacting targets and a broad set of possible true but weakly interacting targets.

**Figure 4:**
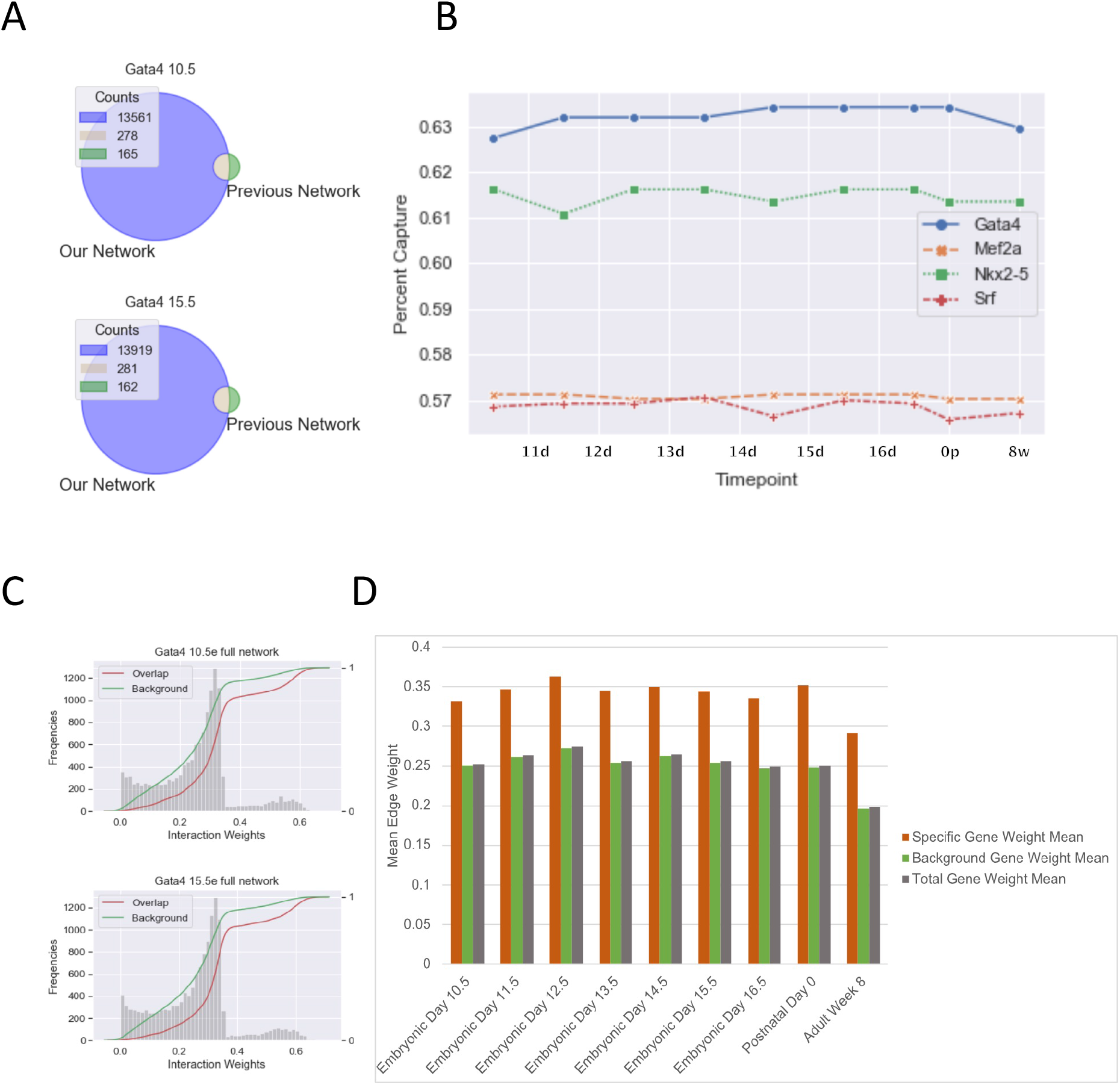
Model Comparison. A) TF-gene interaction Venn Diagram overlap between TReNCo model and previous study B) Line plot showing percent TF-gene interactions captured from a previous study with the TReNCo model C) CDF plot with weight of overlapping interactions vs background in TReNCo model D) Bar plot of mean TF-gene weights for Specific/overlapping interactions (red), all other background interactons (green) and all interactions (blue)

To further support the biological relevance of our networks, we selected the full TF network, the overlapping network, and the connections unique to our network, and performed GO-term enrichment analysis (Figure 5, S9). We see that in the case of Mef2a, there is a similar core of regulatory processes that are maintained from 10.5e and 8w (Figure 5). Of interest, we found that in younger 10.5e hearts there was significant enrichment for development related genes as opposed to older 8w hearts which had T cell activation terms enriched. This makes sense when considering recent studies showing Mef2a involvement in inflammation-related processes and the interaction of T cell activation and inflammation (Skapenko et al. 2005; Xiong et al. 2019). Furthermore, we found that overlapping targets between our data and previous data contained terms enriched for cardiac development while full and unique networks showed enrichment for cardiac related and general biological terms. Thus, it is reasonable to conclude that our network contains true biologically relevant interactions in cardiac tissue throughout development.

**Figure 5:**
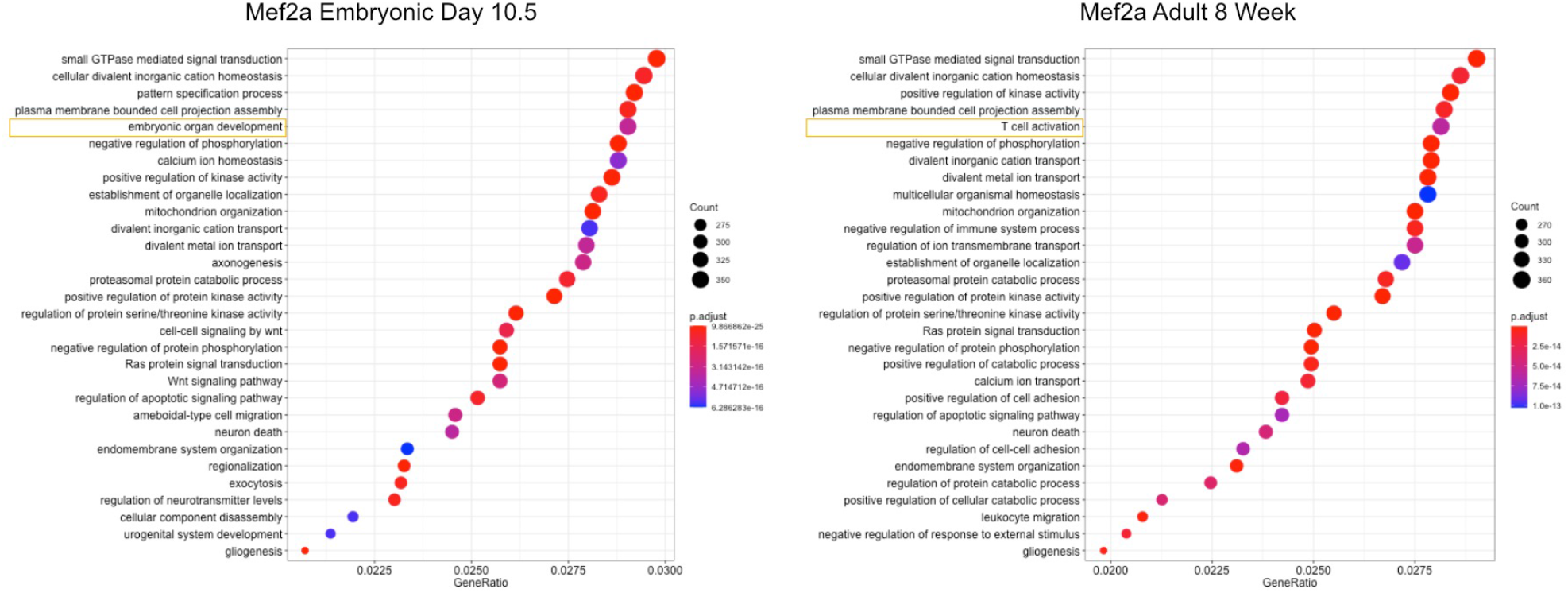
GO-term Enrichment. Enrichment for gene terms between embryonic day 10.5 and 8 week old adult heart. Highlighted terms show first difference in term list

## Conclusions

Our results show that we were able to expand current methods for generating regulatory networks to take advantage of Topologically Associated Domains (TADs) to limit the predicted influence of enhancers. In this way, we were able to produce highly similar results as reported previously with the added benefit of the networks containing an expanded set of potentially relevant biological connections that can be explored. Additionally, we have developed a framework that can be exploited for a diverse array of species and cell types requiring only two experimental assays, H3K27ac ChIP-seq and RNA-seq. We believe this method opens the possibility for understanding deeper connections and new possibilities for biological discovery.

## Materials and Methods

### Data Preprocessing

TReNCo begins by generating distinct transcription start sites (TSSs), using a parsing tool, MakeGencodeTSS, for protein-coding genes from Gencode annotation files: Mouse Gencode version 4 by default. Promoters are then constructed using bedtools slot to extend 1000 nucleotides upstream and 200 nucleotides downstream of the TSS in a strand specific extension. Enhancer boundaries are then generated by using bedops merge to merge the user defined H3K27ac ChIP peak bed files and excluding overlaps with promoter regions.

A transcript expression matrix with normalized log2 TPM is generated from the provided RNA-seq expression tables with each row corresponding to a gene and the columns corresponding to a sample. The same is done for enhancers with bedtools coverage being used to calculate the coverage for each enhancer for each sample using the enhancer regions defined previously.

Log-Odds ratio for TF (TF) binding to promoters and enhancers is calculated with MEME-suite software, FIMO, using cis-bp motif database on promoter and enhancer sequences extracted from the genome, default mm10 (GRCm38), using bedtools getfasta and the bed files generated previously. TF matrices are constructed by reformatting the native output of FIMO and converting TF names to gene symbols

### Model Algorithm and Matrix Factorization

We use the following annotations: M for a matrix, **m**_j_ for a matrix column, and m_i,j_ for a matrix element. TReNCo will, for each sample *k*, extract gene expression (**g**^k^) and enhancer expression (**e**^k^) vectors from the previously built gene (G) and enhancer (E) TPM matrices. From the gene expression vector, TF genes will be extracted from the gene matrix (G) and scaled from 0 to 1 to form a network weighting score (*G’*^*k*^).

Then for each TAD *t* genome interval defined in a bed file, default is provided for mice from a previous publication (Gittenberger-De Groot et al. 2005). Genes, promoters, and enhancers are isolated along with TF scores for promoters and enhancers. A distance weight matrix for the TAD (D^t^) is built between all genes and enhancers and used to build a TAD specific enhancer by gene weight (W^k,t^).

Calculate D^**t**^ distance weights using genomic positions ep_i_ and gp_j_ for the corresponding enhancer start peak position and gene start position.

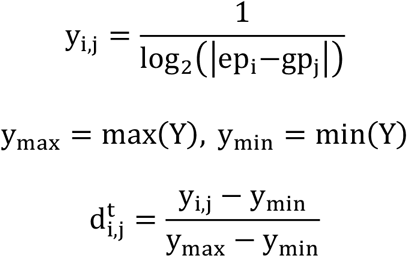

Calculate w^**k,t**^_**i,j**_ enhancer by gene weight

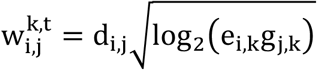

TAD specific TF by enhancer score (M^t^) and TF by gene score (P^t^) are extracted from the FIMO matrices and scaled to the max value of the original matrices. We then factor the vectors and matrices, scale values with an arctan to condense higher values to a similar score approaching 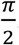. The resulting matrix is weighted by the network weighting score G’^k^ to form TAD specific TF gene regulatory network (R^k,t^).

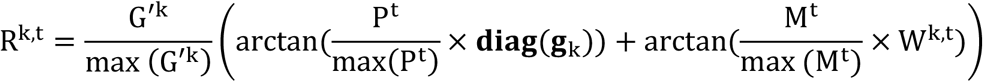

All R^k,t^ matrices are concatenated together to form a full sample specific gene regulatory network R^k^. Gene expression vectors for each sample are converted to gene symbols and output as node weights. The edge matrix is sparse and thus is converted to a vector with TF-gene links being preserved in a string key and zero values being removed. This edge vector is output to a file as the nodes are.

### ENCODE Data

Gene expression and H3K27ac ChIP data was selected from ENCODE. Data was chosen using a script to select mouse heart data corresponding to embryonic day 10.5, 11.5, 12.5, 13.5, 14.5, 15.5, 16.5, postnatal day 0 and 8 weeks old and had a matched set of Gene expression and ChIP-seq. In total every sample had at least technical duplicates with embryonic day 14.5, postnatal day 0, and 8 weeks old time points having 4 replicates of gene expression and embryonic day 14.5 having 4 ChIP-seq replicates. Two of the embryonic day 14.5 gene expression data were dropped due to poor correlation (average R2 less than 0.7) with the rest of the data set. (Table S2 for accession numbers)

### Statistical Analysis and Data Visualization

Plots and graphs were built using seaborn for python and ggplot2 for R scripts. All heatmaps were built in python and analyzed with scikit-learn. Gene networks from previous studies were downloaded from the corresponding journals and converted to a list. Genes networks for Gata4, Srf, Mef2a, and Nfx2-5 were subset from our networks and targets for these TFs were converted to lists. Venn-diagrams of gene list overlaps were built in python using venn2 package. Analysis of GO-terms was performed in R using enrichGO in clusterProfiler with org.Mm.eg.db database. Plots for GO-terms were built using the built in dotplot function in clusterProfiler.

---

**Supplement 1:**
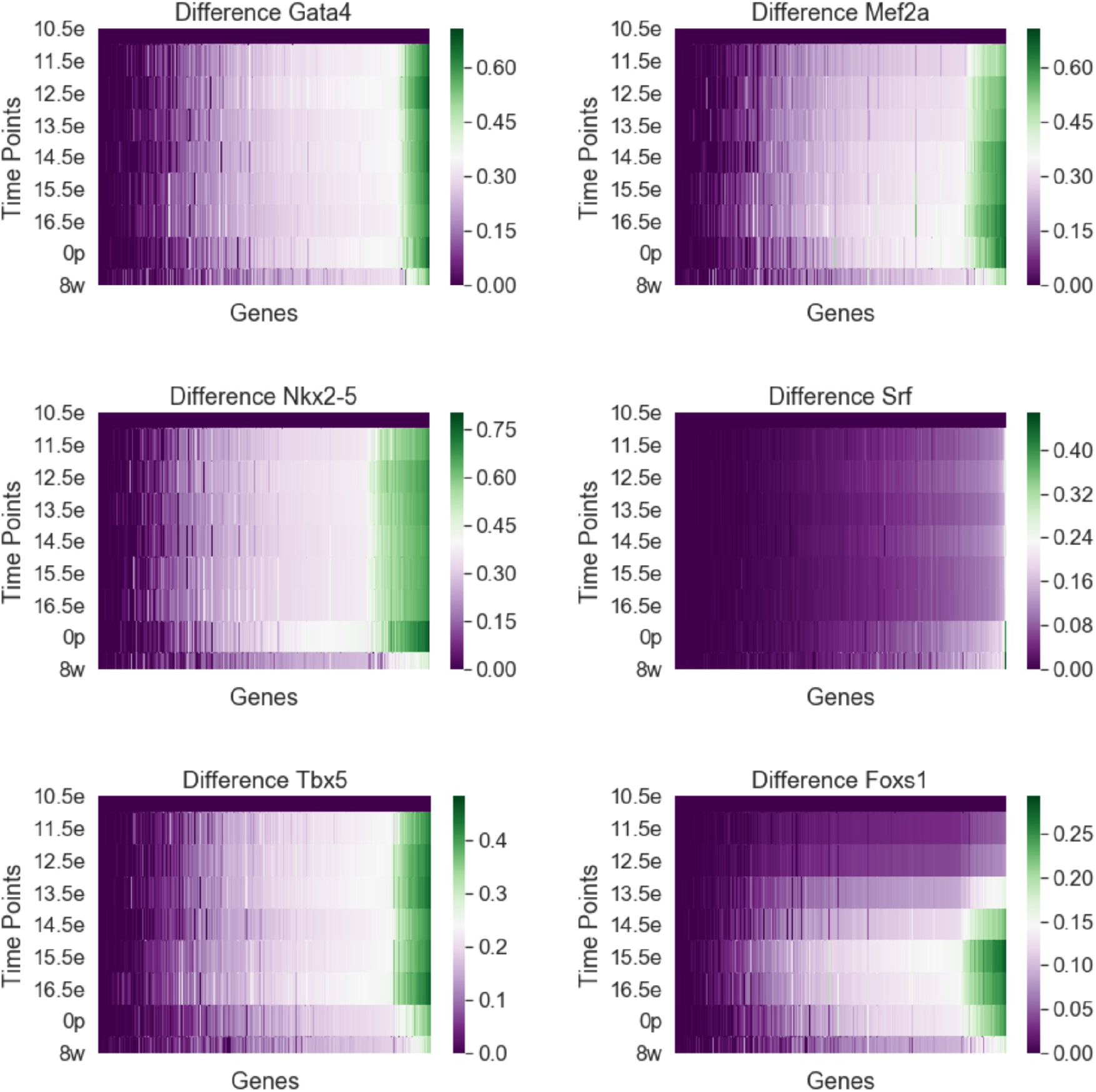
Differential weights

**Supplement 2:**
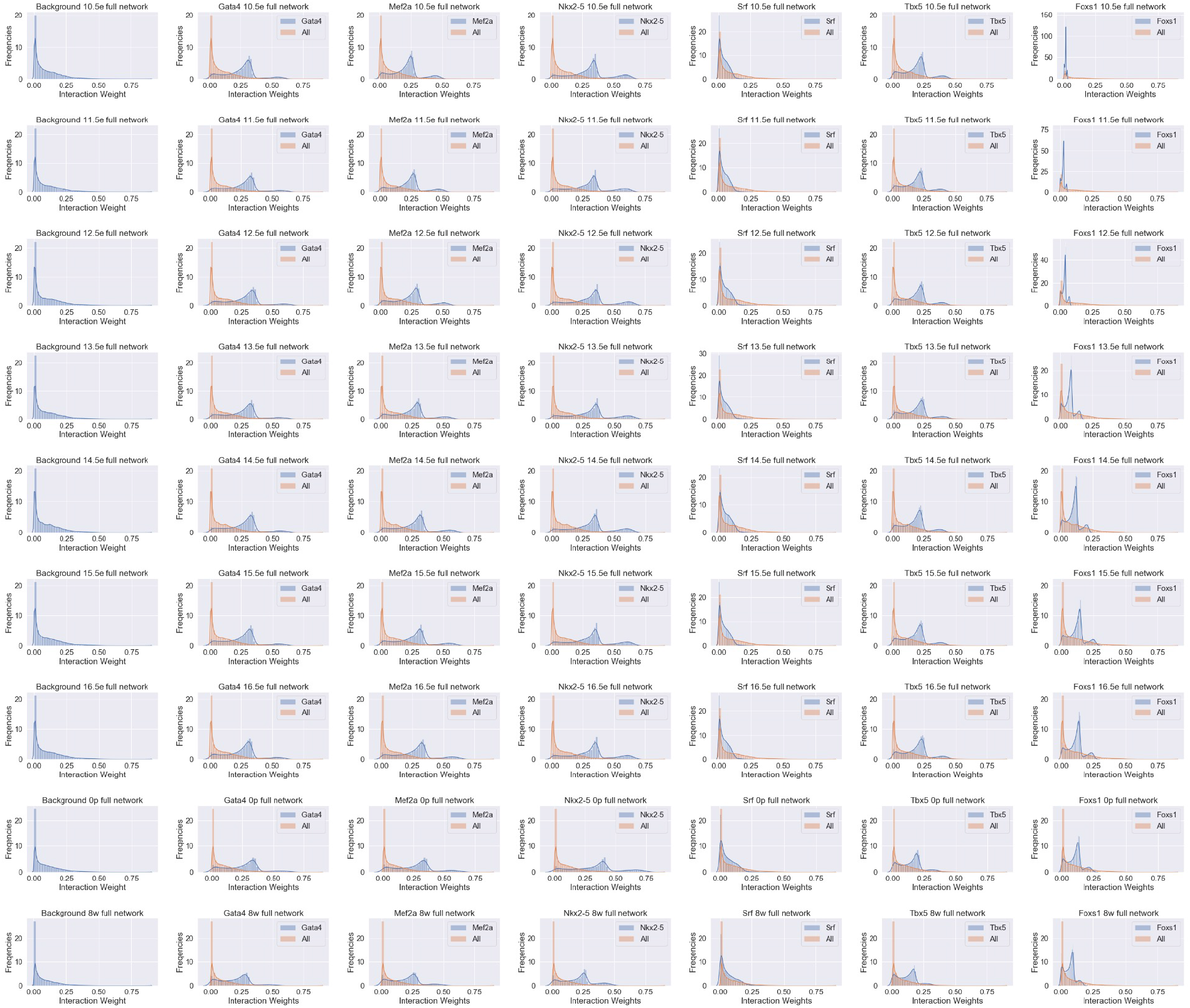
All Distributions

**Supplement 3:**
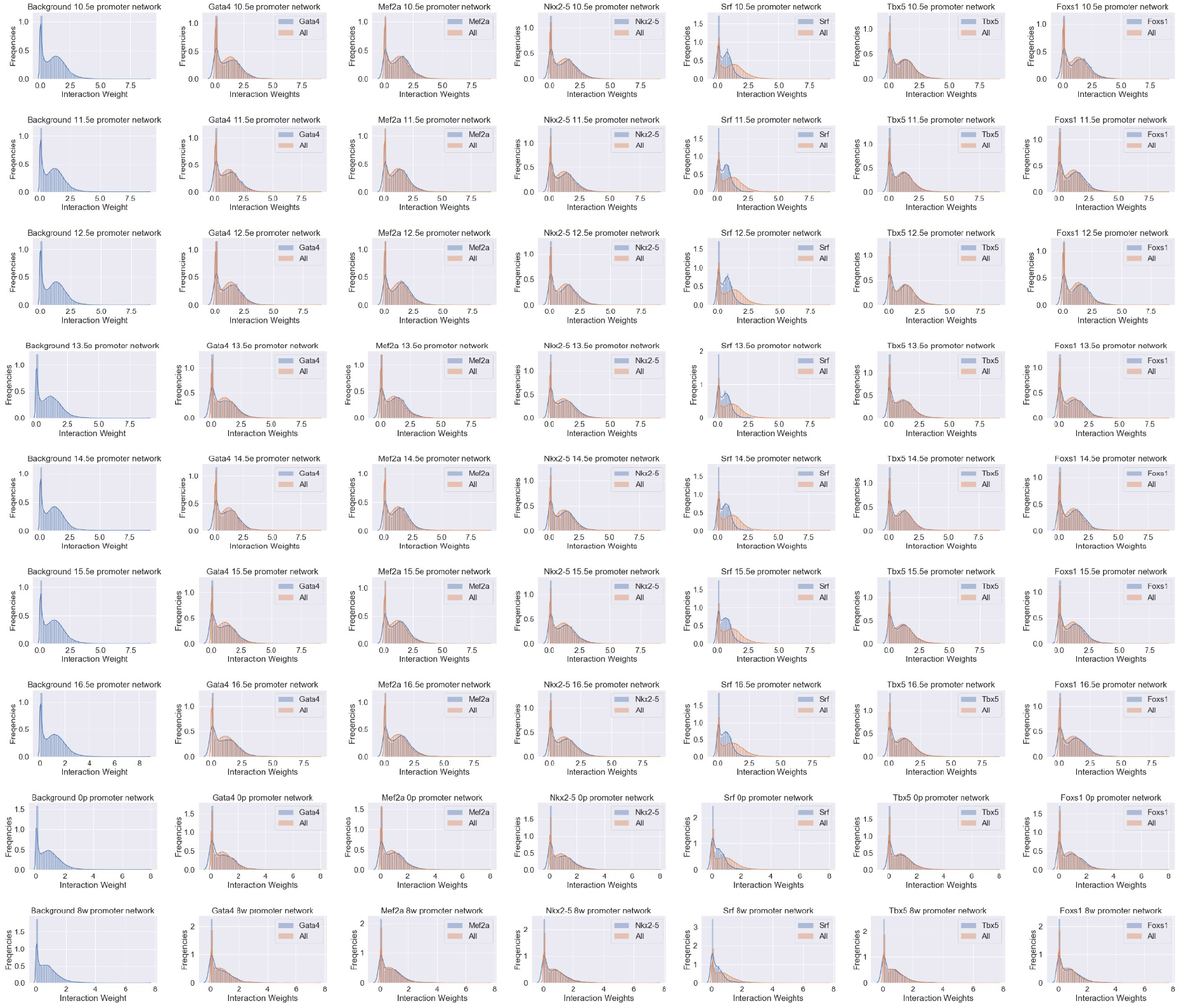
Promoter Distributions

**Supplement 4:**
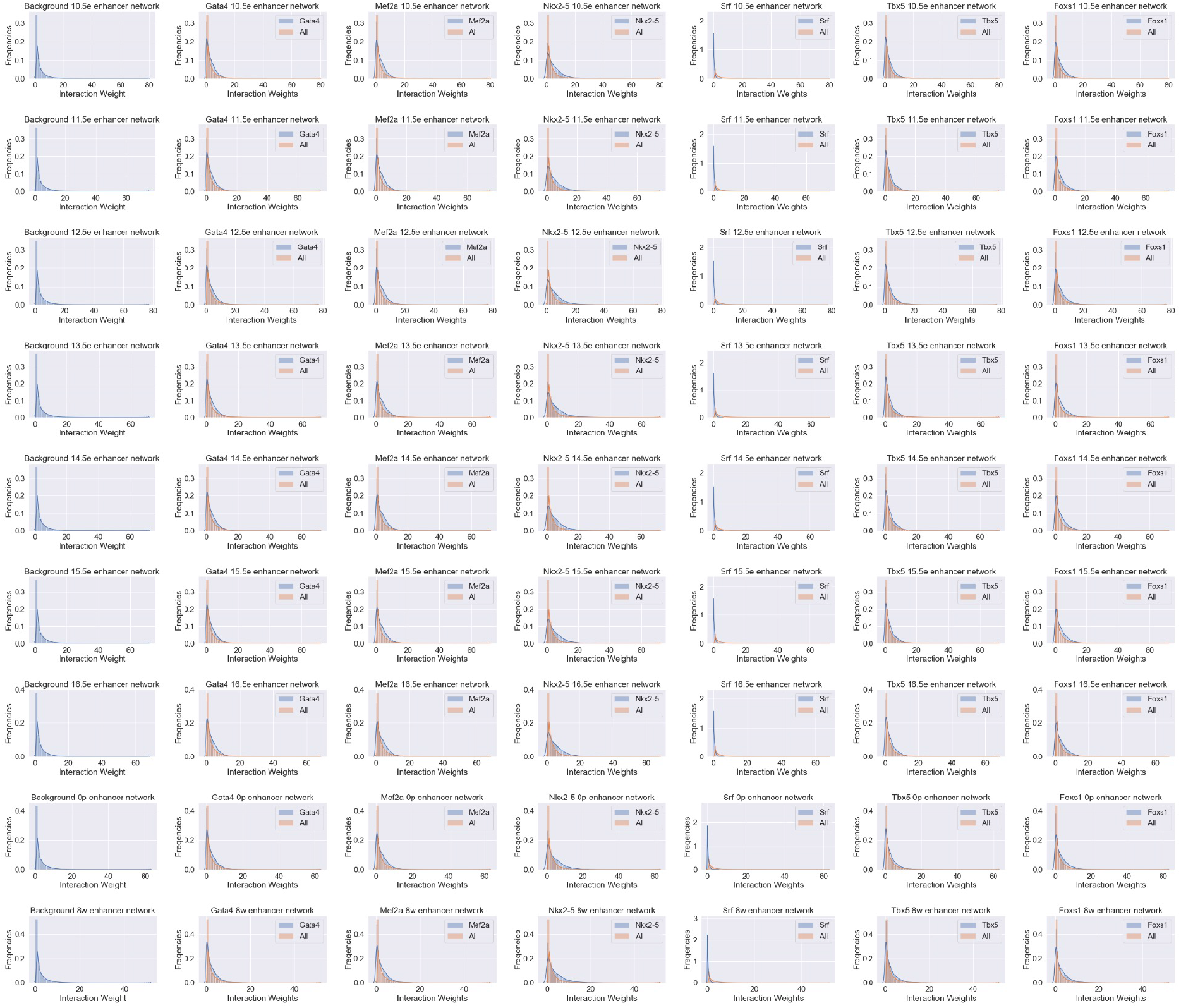
Enhancer Distributions

**Supplemental 5:**
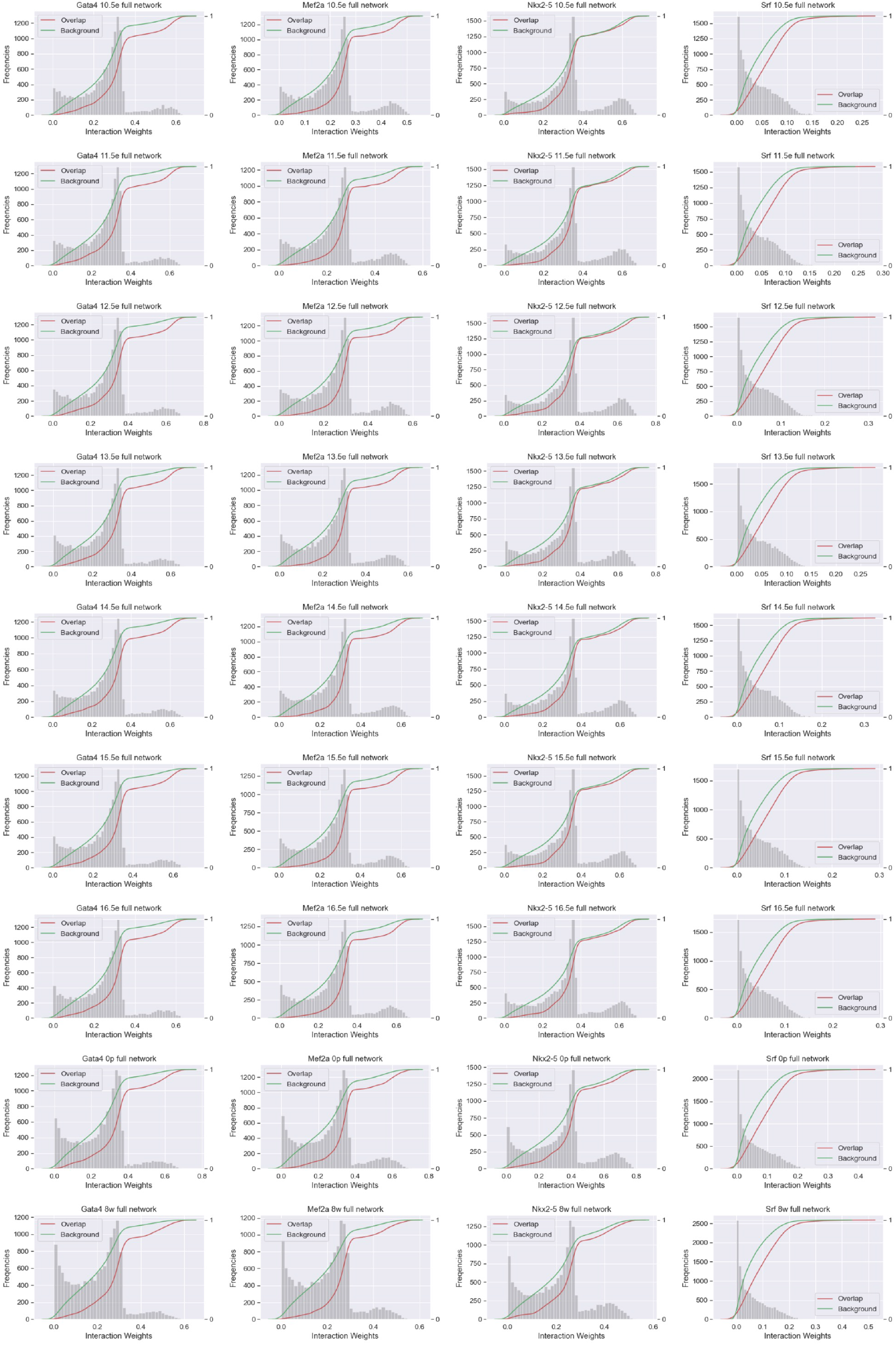
CDF Plots All Genes

**Supplement 6:**
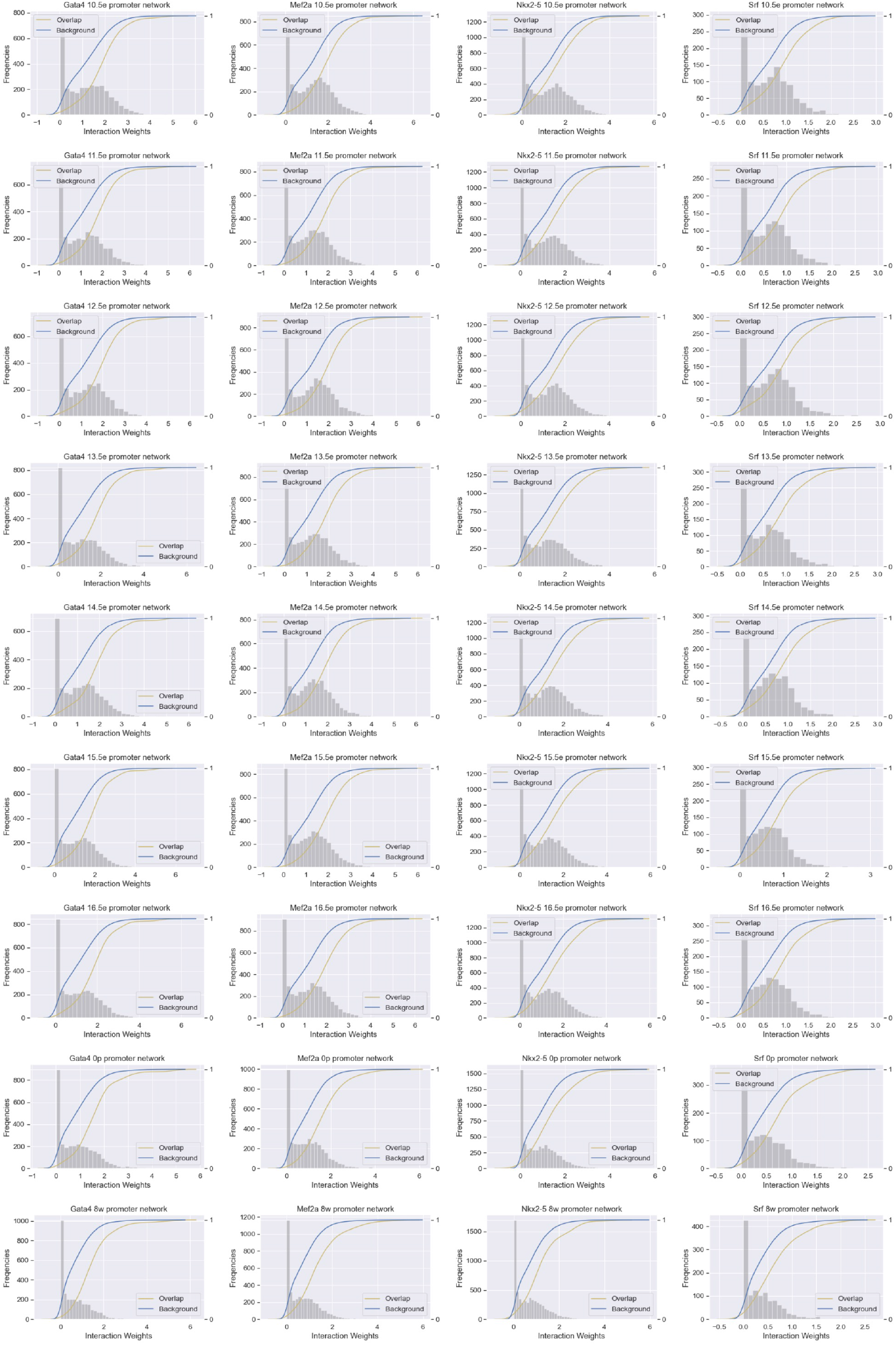
CDF Plots Promoters

**Supplement 7:**
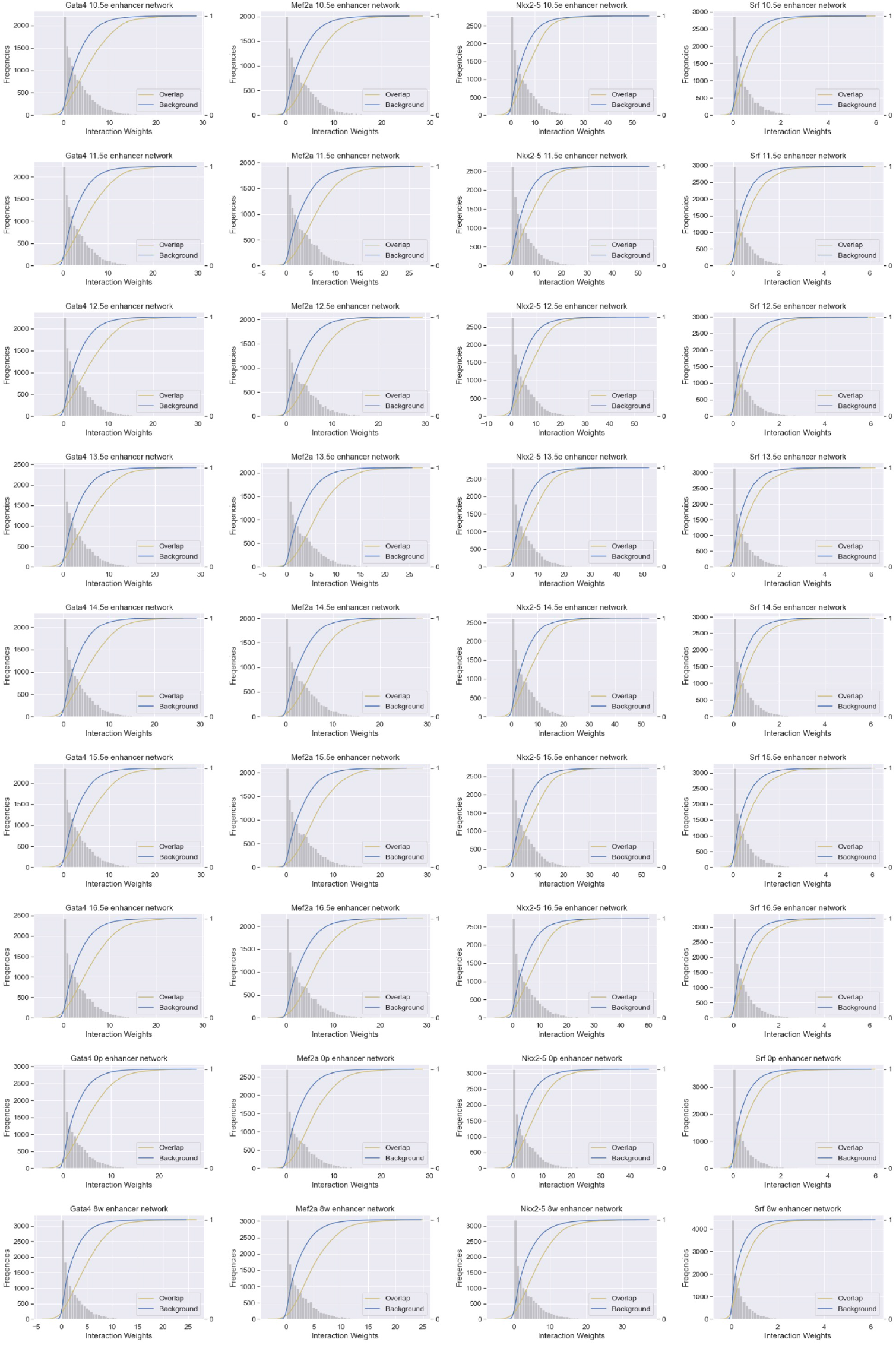
CDF Plots Enhancer

**Supplement 8:**
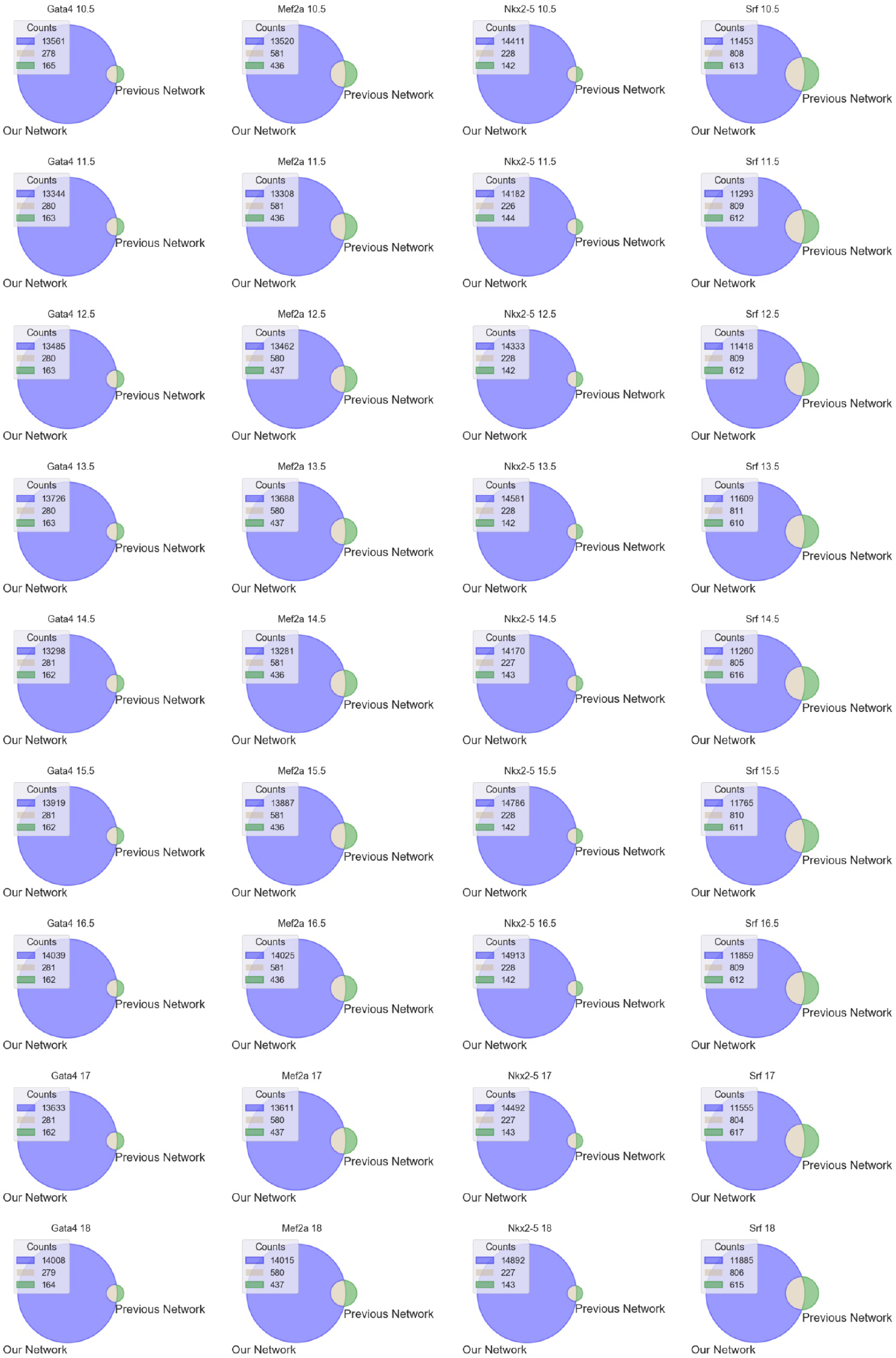
All overlaps

**Supplement 9:**
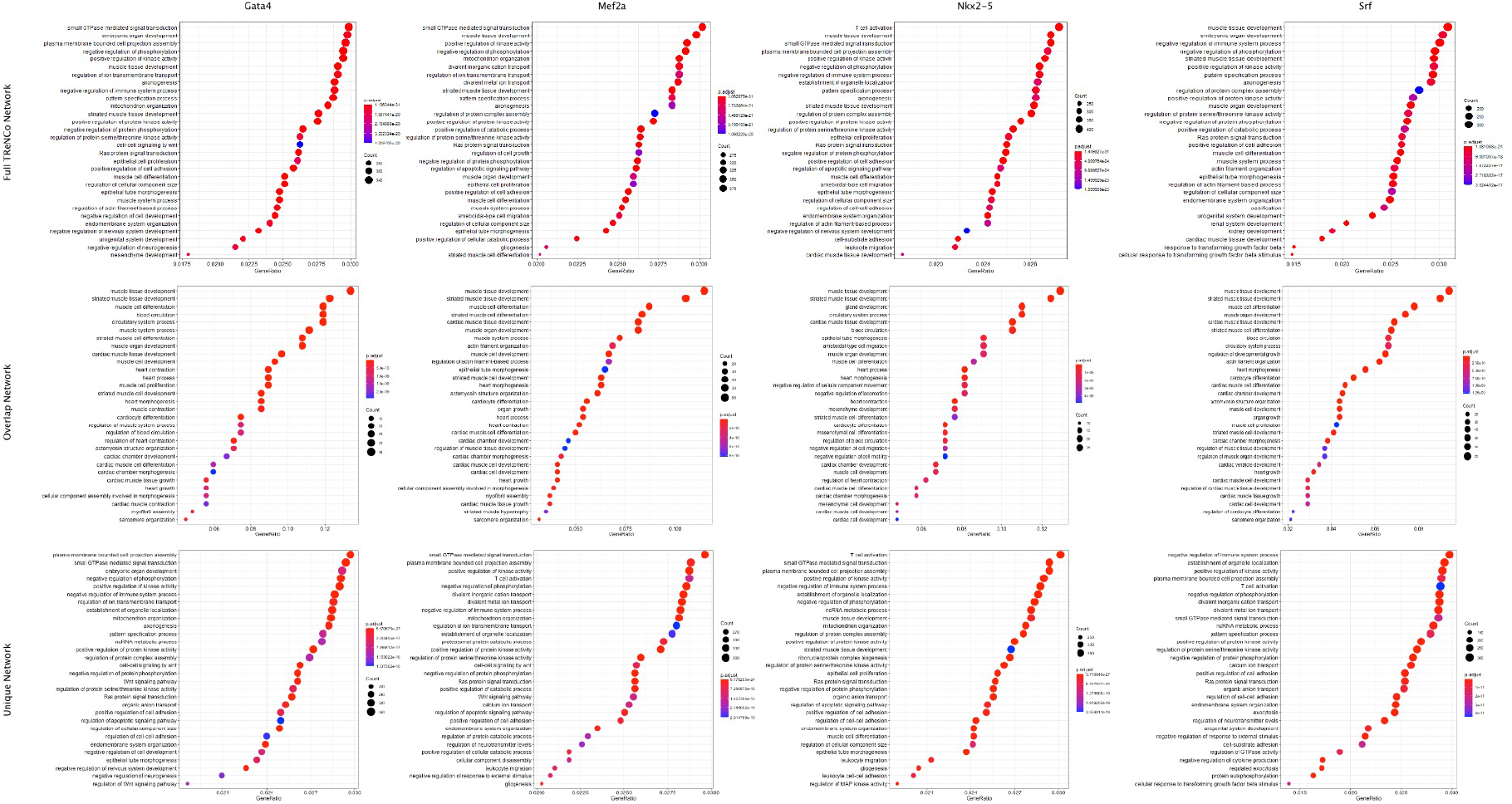
More GO-terms

**Table 1:**
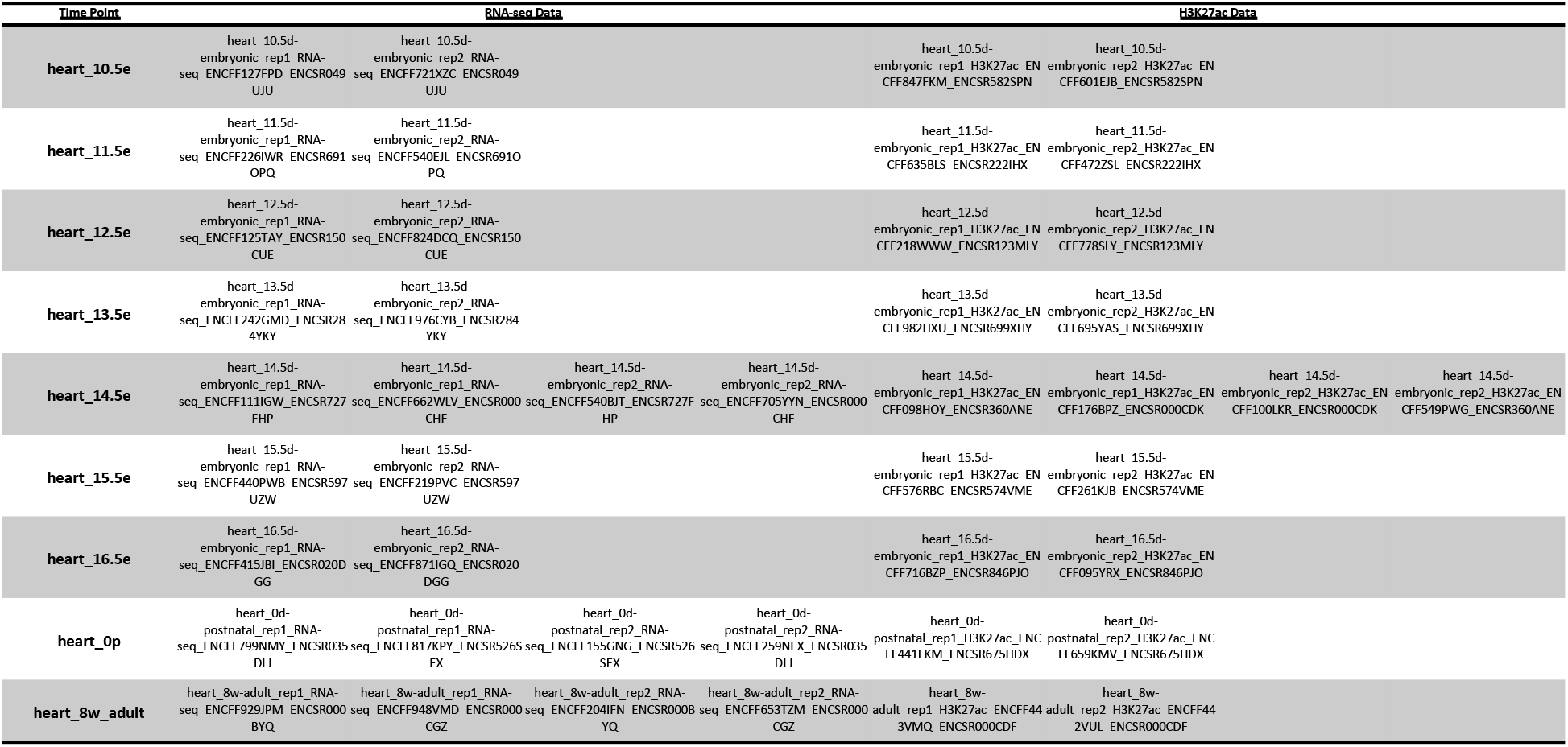
Encode Data for validation

**Table 2:**
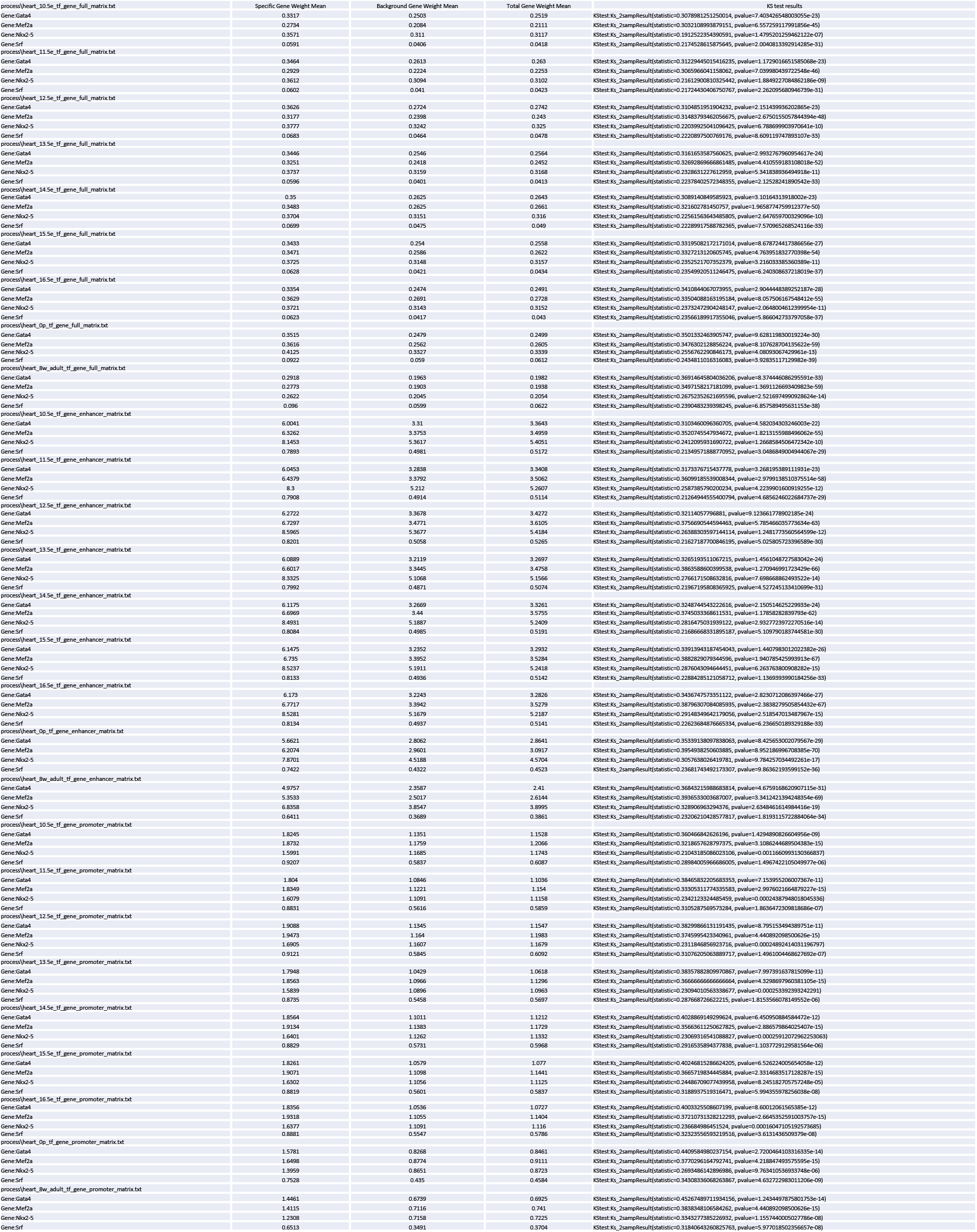
KS-test for all CDFs

## Notes

https://git.biohpc.swmed.edu/BICF/Software/trenco

### Competing Interest Statement

The authors have declared no competing interest.

